# Representation of visual uniformity in the lateral prefrontal cortex

**DOI:** 10.1101/2024.07.07.602149

**Authors:** Yijun Ge, Vincent Taschereau-Dumouchel, Qi Lin, Ali Moharramipour, Zhouyuan Sun, Hakwan Lau

## Abstract

Visual illusions tend to have early visual cortical correlates. However, this general trend may not apply to our subjective impression of a detailed and uniform visual world, which may be considered illusory given the paucity of peripheral processing. Using a psychophysically calibrated visual illusion, we assessed the patterns of hemodynamic activity in the human brain that distinguished between the illusory percept of uniformity in the periphery (i.e., Gabor patches having identical orientations) from the accurate perception of incoherence. We identified voxel patterns in the lateral prefrontal cortex that predicted perceived uniformity, which could also generalize to scene uniformity in naturalistic movies. Because similar representations of visual uniformity can also be found in the intermediate and late layers of a feedforward convolutional neural network, the perception of uniformity may involve high-level coding of abstract properties of the entire scene as a whole, that is distinct from the filling-in of specific details in early visual areas.

## Introduction

Despite limited spatial resolution and color sensitivity in our peripheral vision (Curcio et al., 1990; Strasburger et al., 2011), we perceive the world as relatively uniform and detailed throughout our entire visual field. How does the brain support this uniform perceptual experience given limited peripheral sensory processing and input?

Numerous studies have provided insights related to the subjective perception of uniformity. One line of research on ensemble statistical perception posits that our visual system can extract summary statistics of a visual scene (Alvarez, 2011; Cant & Xu, 2012; Im et al., 2017; Tark et al., 2021; Whitney & Yamanashi Leib, 2018). Instead of creating detailed representations of individual objects, summarizing their statistical properties can efficiently capture the essence of a cluttered scene. This may partly explain the impression of a coherent and rich perceptual experience.

Relatedly, a study using virtual reality (VR) showed that observers often failed to notice desaturation of color in the periphery (i.e., removal of color) (Cohen et al., 2020). They were frequently unaware of these changes, even when only less than 5% of the visual display was actually in color. This finding suggests that our impression of a rich, colorful visual experience across the entire visual field is largely inaccurate. Other studies using static presentation of desaturated stimuli in the periphery have found similar results (Balas & Sinha, 2007; Okubo & Yokosawa, 2023).

Other evidence also indicates that foveal information seems to be extrapolated toward the peripheral visual field (Stewart et al., 2020), akin to perceptual filling-in observed at gaps in the visual field or at the blind spot (Durgin et al., 1995; Komatsu, 2006; Qian et al., 2017). This extrapolation phenomenon is evident in the uniformity illusion, where participants perceive peripheral stimuli gradually changing to match the content of central stimuli, creating the illusion of a uniform visual display (Otten et al., 2017). In a specific version of the illusion, coherent visual patterns presented in the central visual field, such as a set of Gabor patches with identical orientation, can induce the perception of similarly coherent patterns in the periphery, even though the periphery actually contains random, non-coherent stimuli.

The neural basis of the uniformity illusion remains unclear, as current evidence is mostly indirect. Although Suárez-Pinilla et al. (2018) suggest that primary visual cortex (V1) may not be directly involved, this conclusion is based on behavioral inferences and does not rule out the possibility of V1’s involvement through other mechanisms within the visual cortex. Similarly, evidence supporting lateral prefrontal cortex (LPFC) involvement in uniformity perception is also indirect, and remains speculative so far (Lau, 2022). Furthermore, even if the LPFC plays a role, it may influence early visual areas through feedback mechanisms, which could ultimately constitute the neural basis of the illusion.

To address these unresolved issues, we conducted a study to investigate the neural mechanisms underlying the perception of uniformity, utilizing the uniformity illusion. Given that the illusion takes a few seconds to manifest, we were able to determine the threshold duration for each participant at which the illusion occurs half of the time. We employed fMRI to examine brain activity patterns associated with the illusion, and found evidence supporting the involvement of the LPFC in the perception of uniformity.

## Results

### Threshold determination for uniformity illusion

Participants (N=16) first completed a psychophysics experiment to determine their individual threshold duration for perceiving the uniformity illusion. During each trial, participants were presented with a display with central stimuli consisting of coherent Gabors tilted either 45 degrees clockwise or counterclockwise, while peripheral stimuli consisted of randomly tilted Gabors (see Supplementary Fig.1). The duration of stimulus presentation varied randomly from 1.5 to 10.5 seconds in each trial. Then participants were asked to report their perceived uniformity illusion rated from 1 to 4 (Fig.1a).

**Fig 1.**
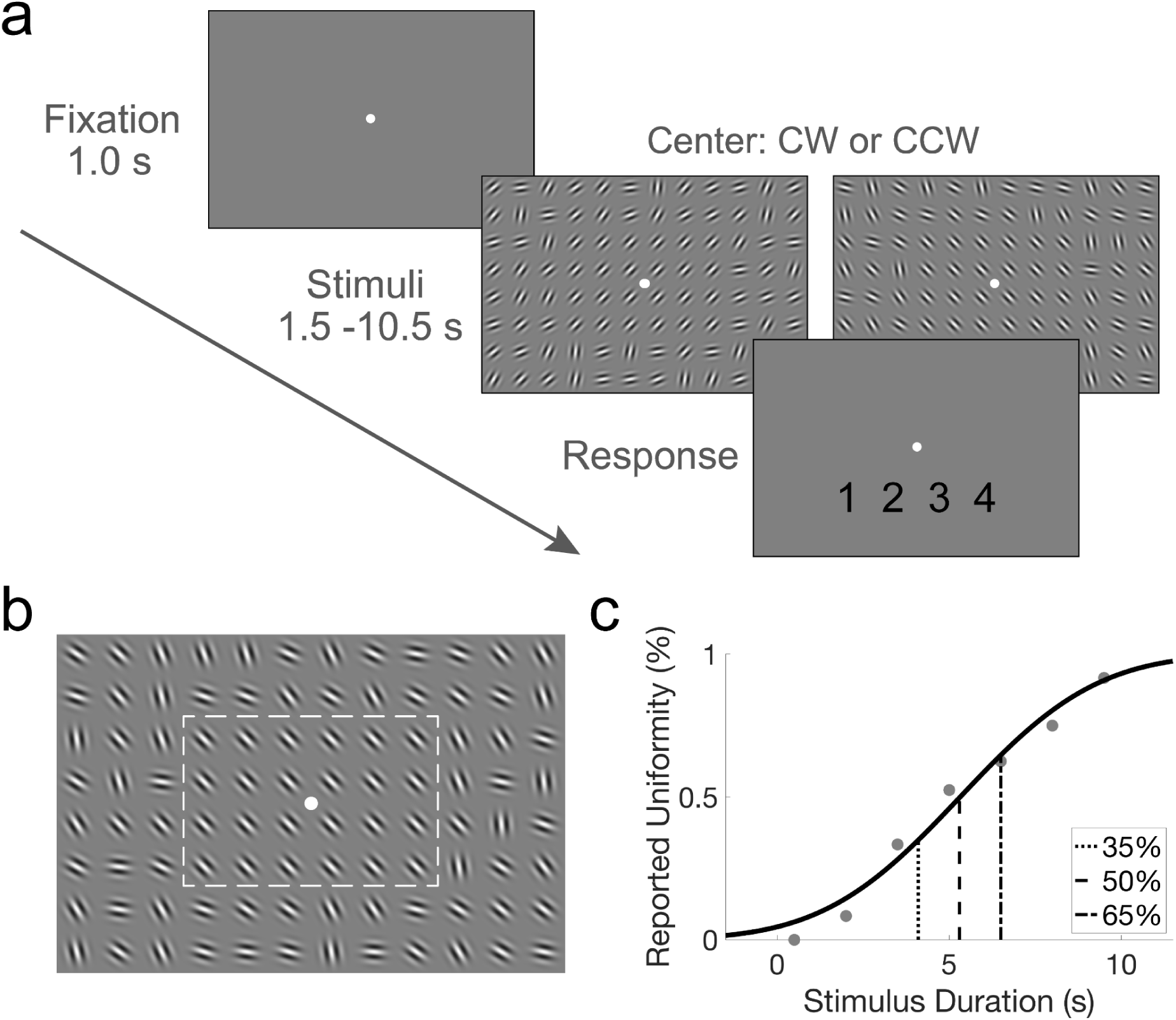
Experimental paradigms and behavior results. **a.** Stimulus presentation sequence for the uniformity illusion threshold duration measurement. Following a 1-second fixation, stimuli were presented for a random duration between 1.5 and 10.5 seconds. The central stimuli comprised coherently tilted Gabors, either clockwise or counterclockwise, while the peripheral stimuli consisted of randomly tilted Gabors. Participants subsequently rated their perceived uniformity illusion on a scale from 1 to 4. **b.** An illustrative example of stimulus, which is not presented to scale here. See supplementary Fig.1 for the actual stimuli used in the experiment, which is more realistically scaled. The dotted line represents the central area of stimuli with coherently tilted Gabors. **c.** The fitted psychometric function with a cumulative Gaussian function for one participant. The threshold durations for perceiving the illusion at 35%, 50%, and 65% levels are indicated. See Supplementary Fig.2 for individual fitted curves and group-level results.

To estimate the threshold duration for perceiving the uniformity illusion at 35%, 50%, and 65% perception levels, a cumulative Gaussian function was fitted to the psychometric data. Fig.1c shows the fitted psychometric function for one participant. The group average threshold durations for perceiving the illusion at 35%, 50% and 65% levels were 2.99 (SD=1.54), 4.74 (SD=1.82), and 6.48 (SD=2.29) seconds, respectively. Individual fitted curves and group-level results are available in Supplementary Fig.2. These tailored threshold durations for each participant were utilized in the following fMRI experiments.

### Decoding the uniformity illusion across the whole brain

In the fMRI experiment, the stimuli and procedure closely resembled those of the psychophysics experiment, except that the stimuli presentation times differed. In half of the trials, stimuli were presented at the 50% threshold duration for each participant. In the remaining half of the trials, stimuli were presented at the 35% and 65% threshold durations, each for an equal number of times. All trials were randomly mixed and presented during each run.

We then performed single-trial General Linear Model (GLM) fitting for the fMRI data, classifying trials into two groups: ‘with-illusion’ and ‘no-illusion’, based on participants’ reports using a median split of ratings ranging from 1 to 4 for each participant (see Supplementary Fig.3 for the response distribution). For this decoding analysis, we focused on trials with 50% threshold duration to ensure consistency in stimulus duration across the analysis. The remaining trials with 35% and 65% threshold durations were utilized in the subsequent generalization analysis. This approach would minimize the influence of a potential confounding factor of stimulus duration when decoding ‘with-illusion’ and ‘no-illusion’ trials.

We conducted decoding analyses across different regions of interest (ROIs) in the brain to investigate the neural representation of the uniformity illusion. ROIs were defined according to the multimodal parcellation of the human cerebral cortex by Glasser et al. (2016) to ensure comprehensive coverage of the whole brain, including early visual regions, higher visual regions, lateral frontal regions, and control auditory regions, among others (see Supplementary Fig.4 and Supplementary Table.1 for the ROI definitions). A Linear Support Vector Machine (SVM) classifier was trained within each ROI to differentiate between trials with and without uniformity illusion. The classifier performance was determined using the Area Under the Curve (AUC) score in a stratified 5-fold cross-validation procedure (regularization parameter C=1), pooling trials across runs. To assess the statistical significance of decoding results, permutation tests were employed. Labels in the test set data were randomly shuffled 1,000 times to generate a null distribution of AUC scores. Additionally, we conducted complementary permutation tests by shuffling all labels and refitting new models 1,000 times (see Supplementary Fig.5a for similar results). To account for systematic differences in signal-to-noise ratio (SNR) and biases in the signals across the ROIs and participants, instead of reporting the raw classification accuracy, we calculated a standardized z-score as the measure of model performance. Z scores were calculated by subtracting the mean of the null distribution (estimated with the permutation test) from the actual AUC and dividing by its standard deviation (Fig.2a). Additionally, to assess decoding significance across participants, we utilized a bootstrap procedure with 10,000 resamples. This allowed us to estimate a p-value indicating the probability of obtaining a group-level mean Z score equal to or smaller than zero, thus evaluating the likelihood of observing significant decoding results by chance.

**Fig 2.**
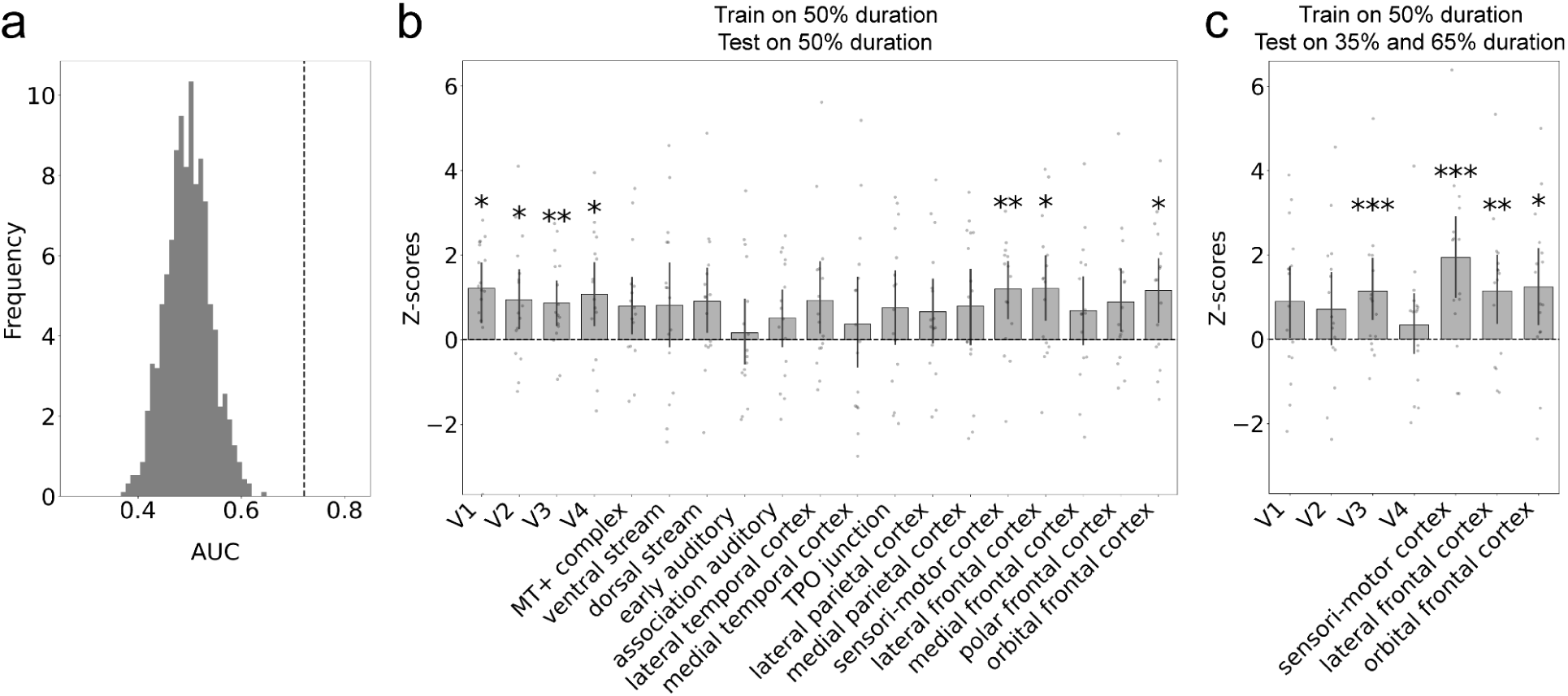
Decoding results using trials with 50% threshold duration and generalization to the trials with 35% and 65% threshold duration. Trials with 50% threshold duration were selected and divided into two groups, with versus without uniformity illusion, based on participants’ reports. A linear SVM classifier was trained on each ROI across the whole brain to classify illusion versus no illusion, yielding AUC scores through 5-fold cross-validation. Permutation tests were conducted by shuffling the labels of the test set data for 1,000 iterations to generate a random distribution of AUC scores (See supplementary Fig.5 with similar results by shuffling labels of both training and test set data in the permutation test). Then the Z score was computed as the difference between the actual AUC and the mean of the random distribution, divided by the standard deviation of the random distribution. A bootstrap procedure with resampling 10,000 times was employed to assess the likelihood of the group-level mean Z score being smaller or equal to 0. **a.** an example of Z score calculation. The dotted line represents the actual AUC score for one participant in one ROI. The distribution illustrates the permutation results after shuffling labels 1,000 times. **b.** Significant group-level mean Z scores were found in V1-V4, sensori-motor cortex, lateral frontal cortex, and orbitofrontal cortex after correction for multiple comparisons. **c.** The trained linear SVM classifiers on the trials with 50% threshold duration, were then applied to the trials with 35% and 65% threshold duration. Only significant ROIs in previous decoding results were selected in the generalization test. Significant generalization results were found in the V3, sensori-motor cortex, lateral and orbital frontal cortex after correction for multiple comparisons. Error bars indicate 95% confidence interval obtained by the bootstrap method. * P<0.05, ** P<0.01, *** P<0.005 after correction.

After correction for multiple comparisons, significant decoding effects were identified in several brain regions (Fig.2b), including V1 (*M* = 1.22, 95% CI = [0.41, 1.81], *P* = 0.029 corrected), V2 (*M* = 0.95, 95% CI = [0.28, 1.65], *P* = 0.024 corrected), V3 (*M* = 0.87, 95% CI = [0.35, 1.38], *P* = 0.006 corrected), V4 (*M* = 1.08, 95% CI = [0.33, 1.80], *P* = 0.036 corrected), sensorimotor cortex (*M* = 1.20, 95% CI = [0.51, 1.84], *P* = 0.007 corrected), lateral frontal cortex (*M* = 1.21, 95% CI = [0.48, 1.97], *P* = 0.013 corrected), and orbitofrontal cortex (*M* = 1.16, 95% CI = [0.42, 1.90], *P* = 0.010 corrected). These findings suggest that distinct brain regions contribute to the illusion of uniformity.

### Generalization analysis of short and long duration trials

Further, we examined whether the linear SVM classifiers trained on trials with 50% threshold duration could effectively distinguish between neural responses indicative of the uniformity illusion in trials with shorter (35%) and longer (65%) threshold durations (see Supplementary Fig.3 for the response distribution). To ensure the specificity of our analysis, only the significant ROIs identified in the previous decoding results were included.

Our results revealed significant generalization effects in several brain regions, including V3 (*M* = 1.14, 95% CI = [0.46, 1.91], *P* = 0.002 corrected), sensorimotor cortex (*M* = 1.94, 95% CI = [1.00, 2.90], *P* < 0.001 corrected), lateral frontal cortex (*M* = 1.15, 95% CI = [0.36, 2.01], *P* = 0.008 corrected) and orbitofrontal cortex (*M* = 1.25, 95% CI = [0.35, 2.14], *P* = 0.012 corrected) (Fig.2c). These findings suggest that the neural representations associated with the perception of the uniformity illusion generalize to external validation data and across different stimulus durations, highlighting the robustness and consistency of the underlying neural mechanisms.

### Generalization analysis of naturalistic movie uniformity in the LPFC

In an effort to circumvent potential confounding effects associated with report-related activities, such as motor planning and execution (Block, 2019; Odegaard et al., 2017), a generalization analysis was conducted under a naturalistic viewing condition without specific task demands. This analysis aimed to predict visual uniformity in movie frames, reasoning that uniformity could be estimated using the variance in the saliency of movie segments, with high variance indicating low uniformity and low variance indicating high uniformity.

To perform this analysis, fMRI data was collected while participants viewed a 20-minute movie clip from the movie Interstellar. Saliency maps for each movie frame were computed using a commonly adopted saliency model (Itti & Koch, 2001), which highlights perceptual salient regions capturing features like contrast, color, and orientation. These maps were generated for each movie frame sampled every second during the viewing period.

The variance in the saliency map, calculated as the standard deviation of saliency values within each frame, served as a measure of saliency variability. High saliency variability, indicating regions that stand out significantly, leads to a perception of non-uniformity. Conversely, low saliency variability, indicating a more uniform distribution of attention-capturing features, creates a perception of visual uniformity.

As such, if high and low variance can be decoded using a linear SVM classifier trained to predict the uniformity illusion, this could indicate a generalization of the decoder. Fig.3a displays the time course of the standard deviation of the saliency map measured for individual movie frames. The top and bottom 33.3% of the distribution were selected to identify frames with high and low saliency variability, respectively, thus categorizing them into low or high uniformity groups. Examples of frames with high and low variance are presented in Fig.3a (top and bottom panels, respectively) and Fig.3b.

**Fig 3.**
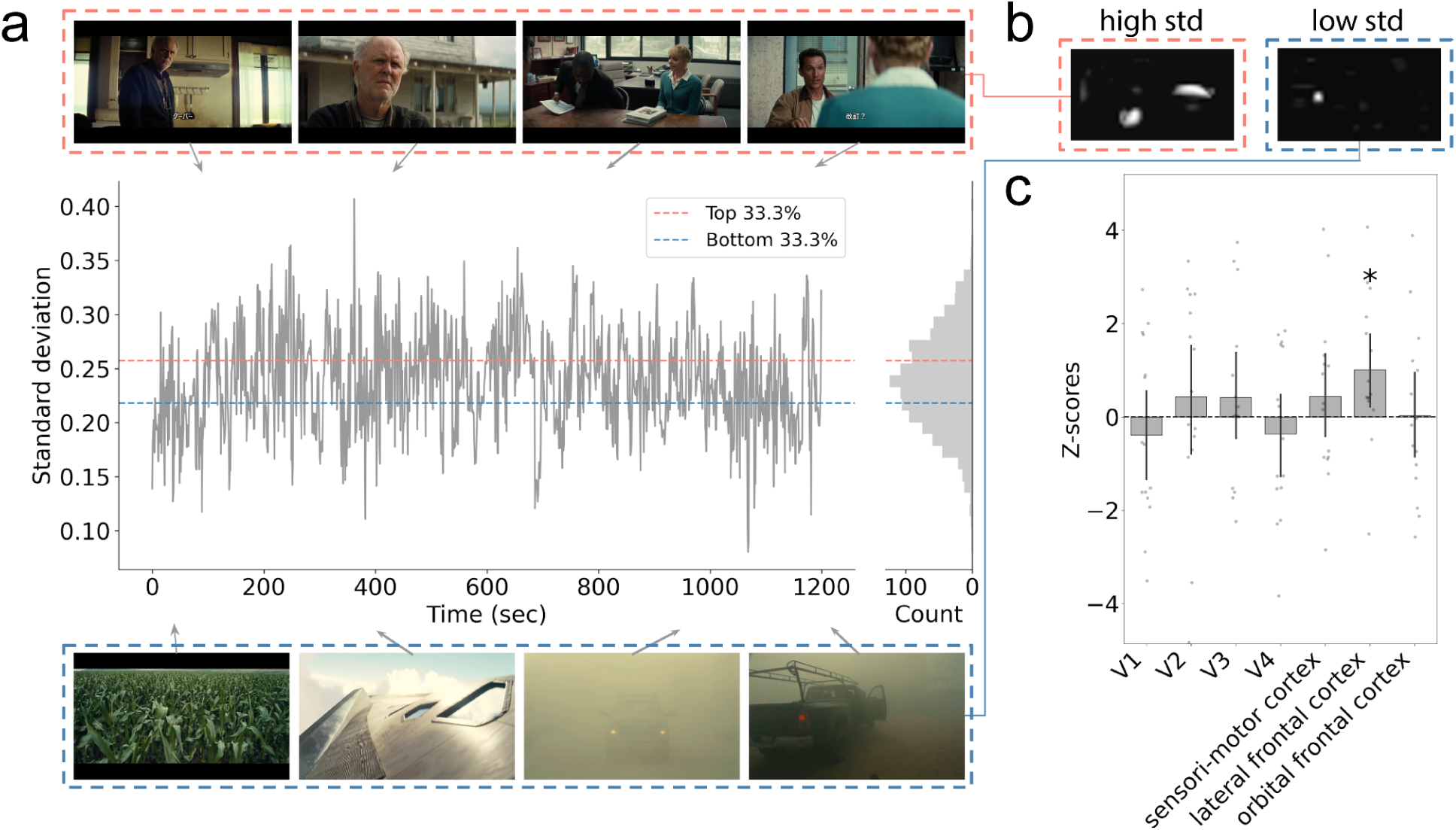
Generalization results on the movie frames. **a.** Participants watched a 20-minute movie clip inside the MRI scanner. Plotted is the time course of the standard deviation of the saliency map measured for the individual movie frames extracted every second. On the right margin, a histogram illustrates the distribution of standard deviation scores across movie frames. The orange and blue dotted lines indicate the top and bottom 33.3% of the distribution, respectively, which delineate movie frames with high and low saliency variability (which in turn means that the frame has low or high uniformity, respectively). The upper (orange) and lower (blue) boxes present examples of movie frames with high and low saliency variability. **b.** Example saliency maps of movie frames with high and low saliency variability. **c.** Generalization results to the fMRI movie data. Movie frames are divided into two groups: frames with the top 33.3% of standard deviation are categorized as ‘Non Uniform’ group, while frames with the bottom 33.3% of standard deviation are categorized as ‘Uniform’ group. The trained linear SVM classifier on the uniformity illusion data (results shown in Fig.2b) was applied to the movie data to predict the saliency variability of each frame. Only the generalization result in the lateral frontal cortex was significant (corrected for multiple comparisons). Error bars indicate 95% confidence intervals obtained by bootstrapping. * P<0.05 after correction.

The generalization analysis revealed significant results only in the lateral frontal cortex (*M* = 1.00, 95% CI = [0.24, 1.77], *P* = 0.042 corrected) (Fig.3c). This indicates that the SVM classifier, trained on uniformity illusion data, was able to predict saliency variability in the movie-watching data. These results underscore the role of the lateral frontal cortex in processing saliency variability across dynamic visual stimuli, thereby substantiating its neural basis for perceiving uniformity in complex, naturalistic settings.

### Distribution of decoding weights in the lateral frontal cortex

We examined the distribution of decoding weights within subregions of the lateral frontal cortex to assess the consistency of neural patterns that predict the perception of visual uniformity. We trained linear SVM classifiers using a feature selection procedure within each fold of cross-validation, as depicted in Fig.2b. Specifically, we selected the top 1200 vertices, the minimum vertex count established across subregions and participants, based on the highest F scores within each ROI.

Fig.4 illustrates the distribution of decoding weights, specifically the selected features, across participants within the subregions of the lateral frontal cortex. For each subregion, we calculated the percentage of participants whose decoding weights covered at least 1% of the selected 1200 vertices. Each subregion is color-coded to reflect this percentage. Darker blue hues represent a higher proportion of participants meeting this criterion, indicating greater consistency of decoding weights across participants within that subregion (see supplementary table.2 for the statistical summaries of decoding weights).

**Fig 4.**
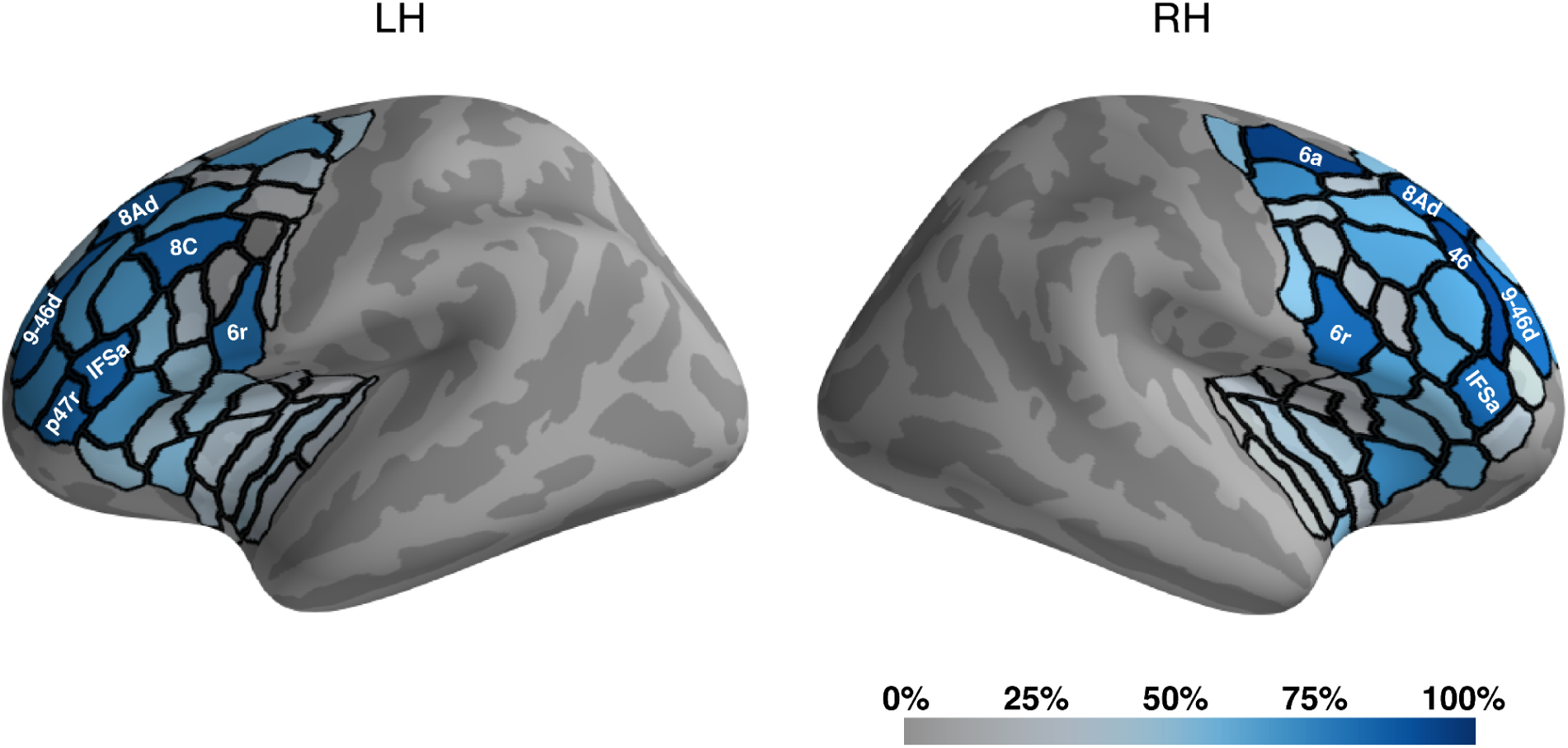
Distribution of the decoding weights in the lateral frontal cortex. The linear SVM classifier was trained on the uniformity illusion data (results shown in Fig.2b) with a feature selection procedure to identify the most informative vertices within each subregion. The distribution of decoding weights within the lateral frontal cortex across participants is depicted, with the subregions color-coded to indicate the consistency of contribution to decoding performance across participants. Darker blue represents a higher proportion of participants showing consistent decoding weights within those subregions, indicating greater consistency across participants. IFSa, 9-46d, 8Ad, and 6r in both hemispheres showed more consistency of decoding weights across participants, with additionally p47r and 8C in the left hemisphere and 6a and 46 in the right hemisphere. See Supplementary Table.2 for the statistical summary for each subregion within the lateral frontal cortex. Anatomical labels are taken from HCP-MMP atlas (Glasser et al. 2016). The results here indicate the scattered distribution of neural patterns which predicted the perception of image uniformity.

We adopted a liberal definition of the lateral frontal cortex ROI, encompassing adjacent areas like the insula, to facilitate a comprehensive exploration of neural representations. This decision was made to enable the SVM classifier to identify significant voxels rather than excluding potentially informative areas *a priori*. Despite this inclusive definition, subsequent analysis reveals that the neural patterns contributing to the decoding of visual uniformity mainly reside within the lateral prefrontal cortex (LPFC).

Our analysis revealed that several subregions of the lateral frontal cortex, including IFSa, 9-46d, 8Ad and 6r in both hemispheres, exhibited more consistent decoding weights across participants. Additionally, regions such as p47r and 8C in the left hemisphere, and 6a and 46 in the right hemisphere, also demonstrated consistency in decoding weights. This scattered distribution of neural patterns within the LPFC suggests a complex and distributed representation underlying the perception of visual uniformity. Notice that this pattern was relatively symmetrical across the hemispheres, suggesting some degree of consistency.

### Decoding image uniformity across CNN layers

Finally, we investigated the representation of image uniformity across different layers of a deep convolutional neural network, ResNet50 (He et al., 2015). The investigation of the biological correspondence of such models has focused mainly on the visual cortex (Khaligh-Razavi & Kriegeskorte, 2014; Yamins et al., 2014). However, our goal here is not to seek a detailed and biological realistic account of our fMRI results. Rather, our modest objective here is just to explore whether even such simple feedforward architecture as ResNet 50 could also support the spontaneous learning of features relevant to uniformity. Fig.5a illustrates the architecture of ResNet50, where 73,000 natural images from the COCO dataset were processed, and principal component analysis (PCA) was applied to extract features explaining 95% of the variance from each of the 50 layers.

**Fig 5.**
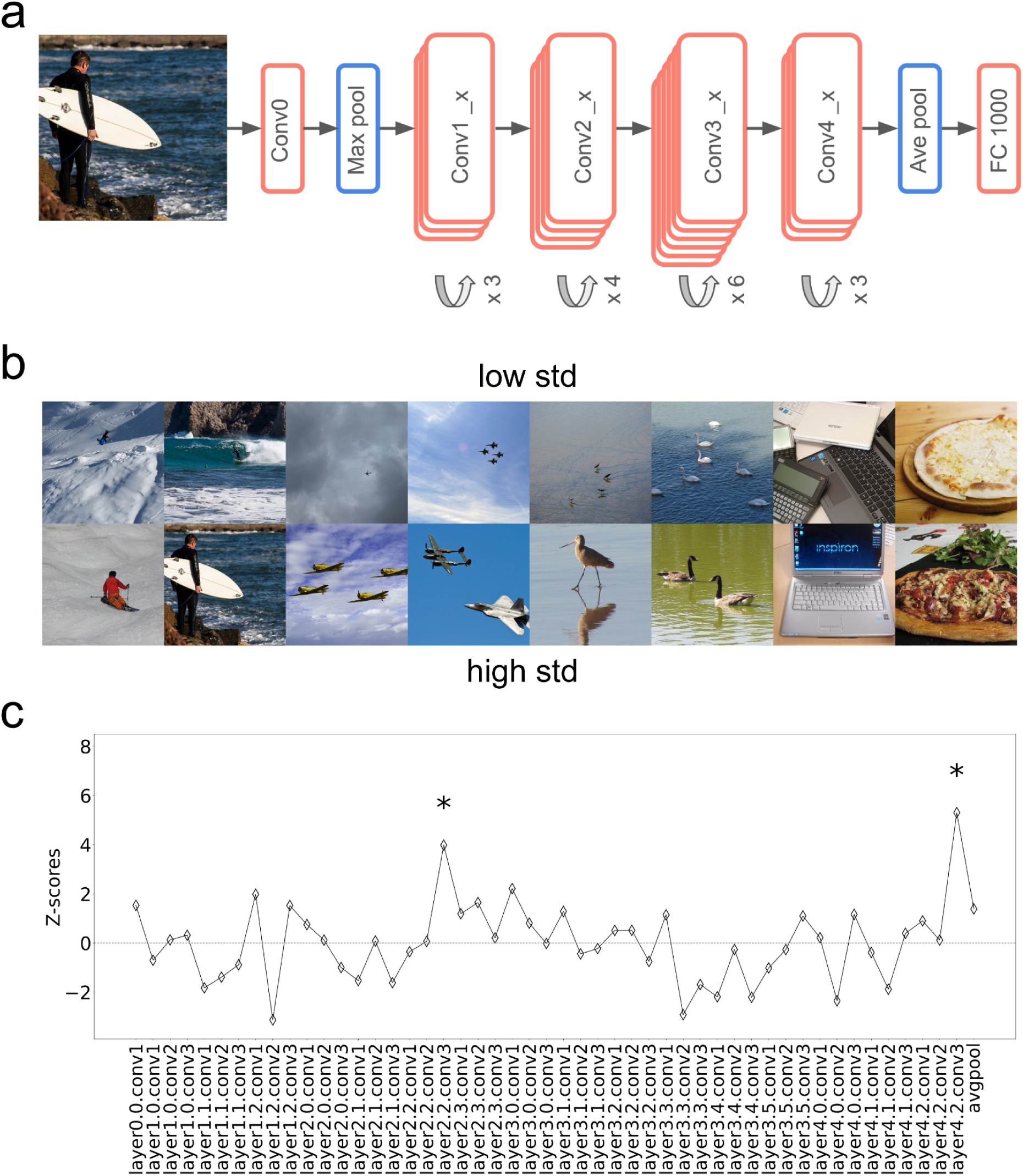
Decoding results for the image uniformity on the deep neural network (ResNet50) layer features. **a.** Schematic of ResNet50 architecture. We fed 73k natural images from the COCO dataset into ResNet50 and applied PCA to extract features that can explain 95% of the variance from each of the 50 layers. **b.** Examples of image pairs with identical semantic labels. To address the confounding issue of semantic labels, only image pairs with identical semantic labels within the top and bottom 33.3% of saliency variability (N=5256 images) were selected. The top and bottom row displays images with low and high standard deviation of saliency map. **c.** Decoding results on the features extracted from ResNet50 layers with 5-fold cross-validation procedure. The linear SVM classifier was trained on layer features extracted from natural images exhibiting high and low saliency variability. The Z scores were obtained from a permutation test,shuffling the test set labels 3,000 times. Significant decoding results were observed in the middle and late layers (19th and 49th) of the ResNet50 after correction for multiple comparisons. This suggests that even in a deep neural network trained for classification of image categories, representations of the image uniformity may already exist at different stages of processing. This observation may parallel the role of the lateral frontal cortex in the visual pathway, as observed in our study (Fig.2-3). * P<0.05 after correction.

To mitigate potential confounding effects of high-level semantic information (e.g, forests tend to be more uniform than indoor scenes), we selected image pairs with identical semantic labels from the top and bottom 33.3% of saliency variability (N=5256 images), as shown in examples provided in Fig.5b. We then trained a linear SVM classifier for each layer features to differentiate between natural images with high and low saliency variability using a stratified 5-fold cross-validation procedure.

To address the issue of varying sizes of different layers, which could lead to overfitting and poor performance, we implemented a feature selection step based on an ANOVA F-test within each fold of cross-validation. This method retained only features with p-values below a stringent threshold (alpha = 0.01), ensuring that only the most informative features were used.

Z scores were obtained from permutation tests by shuffling the test set data 3,000 times to assess the significance of decoding performance (see Supplementary Fig.6 for similar results obtained by shuffling both train and test labels). We opted for 3,000 iterations instead of 1,000 to ensure statistical robustness. This is because we were testing a single model rather than multiple subjects as in the fMRI analysis. After correction for multiple comparisons, we found significant decoding effects in the middle (19th layer: Z=3.98, P=0.033 corrected) and late (49th layer: Z=5.30, P=0.017 corrected) layers of ResNet50 (Fig.5c).

These findings suggest that the representation of image uniformity may be encoded at various stages of processing within a deep neural network, despite the network being primarily trained for object classification in the natural images, rather than recognition of uniformity *per se*. Notably, image uniformity also could not be decoded from early layers. While these results do not directly speak to the biological mechanisms, especially in the prefrontal cortex, we nonetheless think that may provide interesting interpretations of some of our findings, which we will describe in the Discussion section.

## Discussion

In this study, we investigated the neural mechanisms underlying the visual perception of uniformity using psychophysics and multivoxel pattern analysis (MVPA) on fMRI data. We used psychophysics to determine the threshold duration at which each participant experienced the uniformity illusion. The fMRI decoding analyses revealed neural representations of the uniformity illusion in multiple brain regions, including V1, V2, V3, V4, sensorimotor cortex, lateral frontal cortex, and orbitofrontal cortex. Further, we observed that the neural representations in V3, sensorimotor cortex, lateral frontal cortex, and orbitofrontal cortex could generalize to different stimulus durations.

However, it is noteworthy that the generalization to different stimulus durations does not account for potential confounding effects stemming from report-related activity, such as motor planning and execution (Block, 2019; Odegaard et al., 2017). To address this issue, we analyzed data from a naturalistic movie-watching condition, where participants were not given any specific task, and the stimuli were dissimilar to the Gabor patches used in the uniformity illusion. The robust generalization of our decoding results to movie frames with varying levels of uniformity suggests that activity patterns in the lateral prefrontal cortex (LPFC) reflect genuine perceptual processes rather than being solely attributable to report-related responses.

Our findings regarding the involvement of the LPFC in visual uniformity perception are congruent with the view that the prefrontal cortex is causally implicated in high-level visual perception. For example, evidence from Kar and DiCarlo (2021) suggests that recurrent processing from the ventrolateral prefrontal cortex (vlPFC) plays a crucial role in developing behaviorally sufficient object representation. Their study demonstrated that inactivating the vlPFC impairs the late-phase population codes in the inferior temporal (IT) cortex and leads to behavior deficits in object recognition tasks, emphasizing the importance of fast recurrent processing via the vlPFC.

The large receptive fields in the LPFC may be related to its involvement in visual uniformity perception. While the receptive fields in V1 are small and arranged topographically, they increase in size and complexity in higher-order cortical regions (Bentley & Salinas, 2009; Viswanathan & Nieder, 2017). This enlargement and complexity likely aids in processing and integrating diverse visual inputs, possibly influencing the perception of uniformity.

One study on the relationship between the uniformity illusion and affect may provide additional indirect evidence supporting the involvement of higher cortical regions (Dixon et al., 2017; Etkin et al., 2015). The finding indicates that negative emotional states can influence the uniformity illusion by reducing its occurrence and delaying its onset (Kraus et al., 2022). Given that emotional states are more likely to impact higher cognitive mechanisms rather than early visual processes, this implies that the uniformity illusion may be mediated by higher cortical regions such as the LPFC.

Our study may also contribute to existing research on the neural basis of ensemble statistics (Cant & Xu, 2012; Im et al., 2017; Tark et al., 2021), by shedding light on the potential role of the LPFC. Tark et al. (2021) found that ensemble orientations were represented in frontal regions, but these representations were only robust when each mean orientation was linked to a motor response dimension. In our study, however, the generalization of neural patterns in the LPFC to naturalistic stimuli suggests that the involvement of LPFC in processing ensemble statistics may not be solely due to motor response associations.

The concept of subjective inflation (Knotts et al., 2019; Lau, 2022), in which the brain exaggerates sensory input in the periphery, may be relevant to our findings in the LPFC. This inflation account has been distinguished from traditional accounts of perceptual filling-in, which involves the brain completing missing information in detail (e.g., in early visual areas) based on the surrounding context, such as in the blind spot (Durgin et al., 1995; Komatsu, 2006; Qian et al., 2017). In contrast, subjective inflation does not require specific filling-in of information at the sensory level (Odegaard et al., 2018). The presence of robust activation and generalization effects in the LPFC, coupled with the absence of such effects in the early visual areas, appears to provide support for inflation rather than filling-in in relation to uniformity perception.

This interpretation about inflation depends on the null findings in visual areas. Our study found that the neural patterns in the early visual cortex did not generalize well to naturalistic stimuli, suggesting that areas such as V1 may not be the critical region involved in visual uniformity perception. However, this null result, as negative findings in general, especially with methods like fMRI which come with limited sensitivity, should be interpreted with caution. The lack of generalization alone does not definitively rule out the involvement of V1 but rather merely suggests that the evidence is insufficient to support a strong conclusion regarding its role in the perception of uniformity.

Furthermore, the small receptive field size and specific tuning of visual features within the early visual cortex also suggest that it may not be the primary region for perceiving visual uniformity. With a receptive field size of approximately one degree of visual angle (Bentley & Salinas, 2009), the coverage of early visual areas like V1 might be insufficient to integrate information across the entire visual field, which is necessary for generating a sense of uniformity. The uniformity illusion encompasses various visual attributes, including shape, motion, patterns, and identity (Otten et al., 2017). Neurons in the early visual areas primarily respond to simple visual features such as edges, orientations, and spatial frequencies (Grill-Spector & Malach, 2004). This narrow tuning suggests that V1 may not effectively handle the complex integration of diverse visual features required to process the global aspects of visual uniformity.

Additional evidence suggesting that V1 may not be crucial for modulating the uniformity illusion comes from psychophysical studies. Suárez-Pinilla et al. (2018) demonstrated that the tilt aftereffect in the uniformity illusion was influenced by the local physical orientation of the stimulus rather than the global illusory orientation. Given that tilt aftereffect is commonly associated with neural adaptation in V1 (Knapen et al., 2010), their findings suggest that the mechanisms underlying the perception of the uniformity illusion may not involve changes in V1 coding. Had V1 been critically involved, the tilt aftereffect would likely have adapted to the perceived illusory orientation.

Also, in the uniformity illusion, participants seemed able to distinguish between extrapolated patterns in the periphery experienced under the illusion and physically uniform patterns (Otten et al., 2017). This ability to differentiate the conditions may be compatible with the possibility that the illusion of uniformity is not associated with activity in V1 that is exactly identical to stimulation from physically uniform patterns. If the activity patterns were identical as early as in V1, it may be difficult for the subjects to discern them.

Taken together, although our negative results in visual areas are less than conclusive, they may in fact suggest that the mechanismş underlying the uniformity illusion are unlikely to be located within V1, given the congruence with other indirect evidence.

In light of the absence of significant results in early visual areas, our CNN analysis also yields intriguing and consistent findings. Our examination of ResNet50 revealed significant decoding effects in the middle and late layers, which are known for processing higher-level visual features rather than primary sensory inputs. Despite lacking feedback mechanisms, ResNet50 is a simple yet useful feedforward model for visual processing within the visual ventral stream (He et al., 2015; Khaligh-Razavi & Kriegeskorte, 2014; Sexton & Love, 2022; Yamins et al., 2014). Our finding suggests that even for a network trained to perform object recognition rather than perception of uniformity *per se*, representations related to uniformity already emerge. Notably, consistent with our null findings in V1, and the inflation account, such representations are not found in the earliest layers of ResNet50. Although the model is unlikely to directly reflect mechanisms in the prefrontal cortex, the relevant features identified here, especially those in later layers, may be picked up by further downstream processing, which may happen in the prefrontal cortex.

However, if this interpretation is correct, one may expect to find representations of uniformity in areas such as IT as well. One limitation of the current study is that the main findings are driven by training with the stimuli for the uniformity illusion, which differ from the naturalistic video stimuli. Due to the limited amount of data from the videos, they may only suffice for testing generalization, but not necessarily for training the decoders in the first place. Future studies may be needed to address this issue more systematically, by using ample naturalistic movie data for training the fMRI decoders. Also, other network models may also be explored. For example, some recent studies suggest that self-supervised models may provide more realistic explanations of visual processing in the brain (Konkle & Alvarez, 2022; Zhuang et al., 2021).

Another important consideration is that the uniformity illusion typically requires a few seconds to kick in, in contrast to the immediate perception of uniformity often observed with naturalistic stimuli. To address this, future studies could employ paradigms such as the pan-field color illusion (Okubo & Yokosawa, 2023), which induces a more instantaneous effect. This approach may enable a more nuanced comparison of uniformity perception across various contexts, and potentially address mechanisms more directly relevant in naturalistic settings.

Moreover, future research could also explore the involvement of feedforward and feedback mechanisms in the uniformity illusion using techniques such as transcranial magnetic stimulation (TMS) or layer-specific fMRI analysis. TMS can causally probe the contributions of specific brain regions by temporarily disrupting their activity. For instance, the application of theta burst TMS to deactivate the LPFC could assess the causal relationship between LPFC activity and the uniformity illusion. Layer-specific fMRI can distinguish activity in different cortical layers, aiding in the identification of whether perceptual processes are driven by feedforward input to superficial layers or feedback connections to deeper layers. These approaches would provide a more comprehensive understanding of the neural circuits and dynamics underlying perceptual uniformity.

In summary, our study provides new insights into the neural mechanisms underlying the perception of visual uniformity, emphasizing the roles of the LPFC. Our findings suggest that the brain constructs a coherent perceptual experience through complex interactions across multiple levels of the visual processing hierarchy. By utilizing the uniformity illusion as a tool, we have explored the broader phenomenon of how the brain integrates foveal and peripheral information to maintain a consistent and uniform visual experience. Future research should address the limitations of the current study and further elucidate the neural dynamics involved in visual uniformity perception.

## Methods

### Participants

The experiment was approved by the ethics committee of the RIKEN Center for Brain Science. Sixteen healthy participants (4 females, mean age ± standard deviation: 25.4 ± 4.1) were recruited for the behavior and fMRI experiments. All participants completed both the behavior and fMRI experiments, with the exception of one participant who did not complete the movie-watching segment of the fMRI experiment. All participants had normal or corrected-to-normal vision and no history of neurological or psychiatric disorders. Participants were compensated for their participation and provided informed consent in accordance with the guidelines of the Institutional Review Board.

### Psychophysics stimuli and procedures

Stimuli were presented using an MRI-safe projector on a translucent screen with a resolution of 1600 × 900 pixels and a refresh rate of 60 Hz. The experiment was programmed and controlled using PsychoPy (v2002.2.4). Visual stimuli were displayed with visual angles of 32.6 × 18.6 degrees.

The stimuli consisted of a matrix of 32 × 18 Gabors, each with a diameter of 1.02 degrees of visual angle (dva) and a spatial frequency of 4 cycles/degree. The central 12 × 10 Gabors had a coherent orientation tilted 45 degrees clockwise or counterclockwise. The peripheral Gabors had a randomly tilted orientation ranging from 25 to 65 degrees clockwise or counterclockwise (centered around 45 degrees with a uniform distribution within a ± 20 degree range). A fixation dot with a diameter of 0.30 dva was presented at the center of the screen to maintain participants’ gaze.

Prior to the fMRI scanning session, participants underwent one or two practice runs (with half of the trials) outside the scanner environment, lasting approximately ten minutes. This allowed participants to familiarize themselves with the task and experimental procedures. Subsequently, participants completed the main behavior task while lying in the fMRI scanner before scanning. Each trial began with a one-second fixation period. Next, the central stimuli were presented at full contrast, while the peripheral stimuli faded in over 1 second (linear increase in transparency from 0 to 1), facilitating the perception of uniformity illusion by the participants. The stimuli were presented for a randomly chosen duration between 1.5 and 10.5 seconds (including the 1-second fade-in period). Participants were instructed to report their perceived uniformity illusion using a button box held in their left or right hand, with hand assignment counterbalanced across participants. They used four fingers of the designated hand to press buttons numbered 1 through 4, corresponding to their rating of the perceived uniformity illusion (1 = no illusion, 4 = strong illusion). Participants had 2.5 seconds to make their response after the stimulus presentation. Trials without a response within this period were excluded from further analysis.

### fMRI procedures and data acquisition

Participants completed multiple sessions of the experiment during fMRI scanning, with each session lasting between 1 to 1.5 hours. Typically, we aimed to conduct up to three sessions to maximize data collection. But practical considerations such as time constraints or participant availability occasionally necessitated fewer sessions or runs. Eight participants completed three sessions, seven participants completed two sessions, and one participant completed one session.

The fMRI procedure closely mirrored the psychophysics experiment, with stimulus presentation times varying based on each participant’s measured threshold duration. During each run, stimuli were presented in a mixed and randomized order. In half of the trials, stimuli were presented at the 50% threshold duration. In the remaining half of the trials, stimuli were presented at the 35% and 65% threshold durations an equal number of times for each participant. Each run contained the same number of trials, but the run lengths varied due to the variable trial durations between participants. On average, participants completed 8.25 ± 2.77 runs. During the entire fMRI data collection process, participants also underwent a 30-minute retinotopic mapping procedure and viewed a 20-minute movie clip passively.

MRI data were acquired using a Siemens 3T scanner (Prisma) with a 64-channel head coil at RIKEN Center for Brain Science (CBS). Functional images were acquired with a gradient echo planar imaging sequence (2 mm isotropic voxels, 60 axial slices, 96 × 96 × 60 matrix, TR/TE=1000/30 ms, flip angle = 64°, multi-band factor of 4). High-resolution anatomical volume was obtained with a T1-3DMPRAGE sequence (0.7 mm isotropic voxels, 290 × 320 × 256 matrix, TI/TR/TE=1100/2180/2.95 ms, flip angle = 8°).

### Psychophysics data analysis

The psychophysical data were analyzed using custom MATLAB scripts (MathWorks Inc., R2021b). To estimate the threshold duration for perceiving the uniformity illusion, we fitted the psychometric data using a non-linear regression procedure with a cumulative Gaussian function. This fitting procedure allowed us to estimate the duration at which participants perceived the uniformity illusion 50% of the time (as well as 35% and 65%), providing a threshold measurement for the subsequent fMRI experiments.

### fMRI data analysis

The MRI data were analyzed using AFNI (v22.0.11), Freesurfer (v7.2.0), and custom scripts in MATLAB and Python. Preprocessing included B0 field map correction, despiking, slice timing correction, physiological signal correction, and motion correction, with the high-resolution T1 volume co-registered to the minimum outlier volume of the functional images.

A General Linear Model (GLM) was employed to estimate the fMRI response to the uniformity illusion stimulus for each individual trial. Population Receptive Field (pRF) model fitting was performed using the SamSrf Matlab toolbox (version 8.2, https://osf.io/2rgsm/). Visual regions of interests (ROIs) V1 to V4 were manually delineated based on the eccentricity map, polar angle map, and field-sign map, following standard criteria (Infanti & Schwarzkopf, 2020).

For the remaining brain ROIs, we utilized Glasser’s MMP-HCP atlas (Glasser et al., 2016). The HCP atlas was transformed from fsaverage standard space to native surface space for each participant. After excluding the V1-V4 regions, the remaining cortical areas were organized into 15 regions using similar methods as described in Glasser et al.’s paper: (1) dorsal stream; (2) ventral stream; (3) MT+ complex; (4) early auditory cortex; (5) association auditory cortex; (6) lateral temporal cortex; (7) medial temporal cortex; (8) temporal-parietal-occipital (TPO) junction; (9) lateral parietal cortex; (10) medial parietal cortex; (11) sensori-motor cortex; (12) lateral frontal cortex; (13) medial frontal cortex; (14) polar frontal cortex; (15) orbital frontal cortex. Details regarding these ROIs can be found in Supplementary Table.1.

### Decoding analysis

Multivariate pattern analysis (MVPA) was conducted using scikit-learn 1.4.0 (https://scikit-learn.org/stable/). We aimed to discriminate neural patterns between ‘with-illusion’ and ‘no-illusion’ groups using data from trials with 50% threshold duration.

For the MVPA, a stratified 5-fold cross-validation approach was utilized with a linear Support Vector Machine (SVM) classifier. During each fold, feature normalization was applied exclusively to the training data to ensure that the mean and standard deviation were standardized to zero and unity, respectively. This normalization step aimed to standardize the feature space, facilitating fair comparisons and robust classification across vertices with varying feature distributions.

Feature selection was performed on the training data based on F-scores, which compare the variance between the ‘with-illusion’ and ‘no-illusion’ conditions. The F-score quantifies the extent to which a feature discriminates between the two conditions, allowing us to identify the top 1200 vertices. This selection was based on the minimum number of vertices present within an individual ROI across participants. This methodology ensured that informative features were retained while accounting for variability in ROI size across participants, thereby optimizing classification performance. Classification accuracy was evaluated using Area Under the Curve (AUC) scores, with the regularization parameter C set to 1.0.

To assess the statistical significance of the decoding results, permutation tests were employed. The labels in the test set data were randomly shuffled 1,000 times to create a null distribution of AUC scores. Additionally, a complementary permutation test was conducted by shuffling all labels in both the train and test set, and then refitting new models 1,000 times, which yielded similar results. Subsequently, Z scores were calculated by subtracting the mean of the null distribution from the actual AUC and dividing by its standard deviation (see Supplementary Fig.8 for the raw AUC score results, which exhibit similar outcomes). Then, a bootstrap procedure with 10,000 resamples was utilized to estimate the probability of obtaining a group-level mean Z score equal to or smaller than zero. Significant decoding effects were identified following corrections for multiple comparisons using the Holm correction method. The Holm correction (step-down procedure using Bonferroni adjustment) was applied using the ‘multipletests’ function from the statsmodels library in Python with an alpha level of 0.05 (method=’holm’).

Generalization analyses were conducted to apply the SVM classifier, which was trained on trials at the 50% threshold duration, to trials with 35% and 65% threshold durations. This analysis included only the significant ROIs identified in the initial decoding results. Similar permutation tests and a bootstrap procedure were used to assess the statistical significance of the generalization effects.

### Movie-watching fMRI data analysis

Participants viewed a 20-minute clip from the movie *Interstellar* without performing any specific task while their fMRI data were recorded. The same preprocessing procedures described above were applied to the movie-watching fMRI data, excluding the GLM fitting step. Preprocessed fMRI data were delayed by 6 seconds to account for the hemodynamic response, aligning with the movie frame extracted every second. Given our TR of 1 second, this resulted in 1200 movie frames with corresponding fMRI volumes matched to each visual stimulus (every movie frame).

Saliency maps were computed for each movie frame using the Itti & Koch saliency model (Walther & Koch, 2006) implemented in the SaliencyToolbox (available at https://github.com/DirkBWalther/SaliencyToolbox). Saliency maps highlight regions in an image or frame that are perceptually salient to human observers, capturing features such as contrast, color, and orientation that attract attention. The standard deviation of the saliency map was calculated to measure saliency variability. We selected the top and bottom 33.3% of movie frames in the saliency variability distribution to categorize frames into high and low saliency variability groups, respectively.

For the generalization analysis of movie-watching fMRI data, the SVM classifier trained on the uniformity illusion data was used to predict the variability of saliency across different frames in the movie. We identified the fMRI volumes that corresponded to movie frames categorized into groups of high and low saliency variability. We evaluated the generalization results to determine the extent to which neural representations transferred from the uniformity illusion to dynamic visual stimuli in the naturalistic movie. We used similar permutation tests and a bootstrap procedure to test the statistical significance of the generalization effects.

### Decoding analysis in the CNN layer features

We investigated the representation of image uniformity within the features of a deep convolutional neural network, specifically the ResNet50 model. The ResNet50 architecture, comprising 50 layers, was used to process 73,000 natural images from the COCO dataset.

Principal component analysis (PCA) was applied to extract features from each layer, retaining components that accounted for 95% of the variance. This process of dimensionality reduction ensured that the most important features were preserved for subsequent analysis.

To address potential confounding effects related to semantic labels, we selected image pairs with identical semantic labels from the COCO dataset. Specifically, we focused on images within the top and bottom 33.3% of saliency variability, resulting in a subset of 5,256 images. The saliency variability of these images was measured using the Itti & Koch saliency model. Fig.5b provides examples of selected image pairs, illustrating high and low saliency variability within categories that are semantically identical. Supplementary Fig.7 provides additional examples of the semantic labels.

For the decoding analysis, a linear SVM classifier was trained on the features extracted from each of the 50 layers of ResNet50. The objective of the classifier was to distinguish between images with high and low saliency variability. To ensure reliable and unbiased classification performance, a stratified 5-fold cross-validation procedure was employed. In each fold, the data was divided into training and test sets. The training set was normalized to have a mean of zero and a standard deviation of one. For feature selection, we employed a method based on the False Positive Rate (FPR) test to control the rate of false detections on the training data (Tang et al., 2014). Specifically, we utilized an ANOVA F-test to calculate p-values for each feature, selecting those with p-values below a stringent threshold of alpha = 0.01. This ensured that only features with a significant statistical association to the target variable were retained, thereby improving the model’s performance by focusing on the most informative features. To assess the statistical significance of the decoding performance, permutation tests similar to those used in the previous analysis were conducted (see Supplementary Fig.9 for the raw AUC scores, which show similar results). A total of 3,000 iterations were performed, and the p-value was calculated as the proportion of permuted Area Under the Curve (AUC) scores that exceeded the actual AUC score.

## Data availability

The data that support the findings of this study will be made available on OpenNeuro upon acceptance of the manuscript.

## Code availability

Codes will be available at Open Science Framework (https://doi.org/10.17605/OSF.IO/NVJ9G).

## Acknowledgements

This study is supported by internal funding at the RIKEN Center for Brain Science (to H.L.).

## Author contributions

Y.G. and H.L. designed the study; Y.G. performed the experiments; Y.G., Q.L., A.M., and Z.S. analyzed the data under the guidance of H.L.; Y.G., V.T.-D., Q.L., A.M., Z.S., and H.L. wrote and edited the manuscript.

