## Supplementary Information for "Representation of visual uniformity in the lateral prefrontal cortex"

Contents of supplementary information:

Supplementary Figure 1 to 9
Supplementary Table 1 to 2

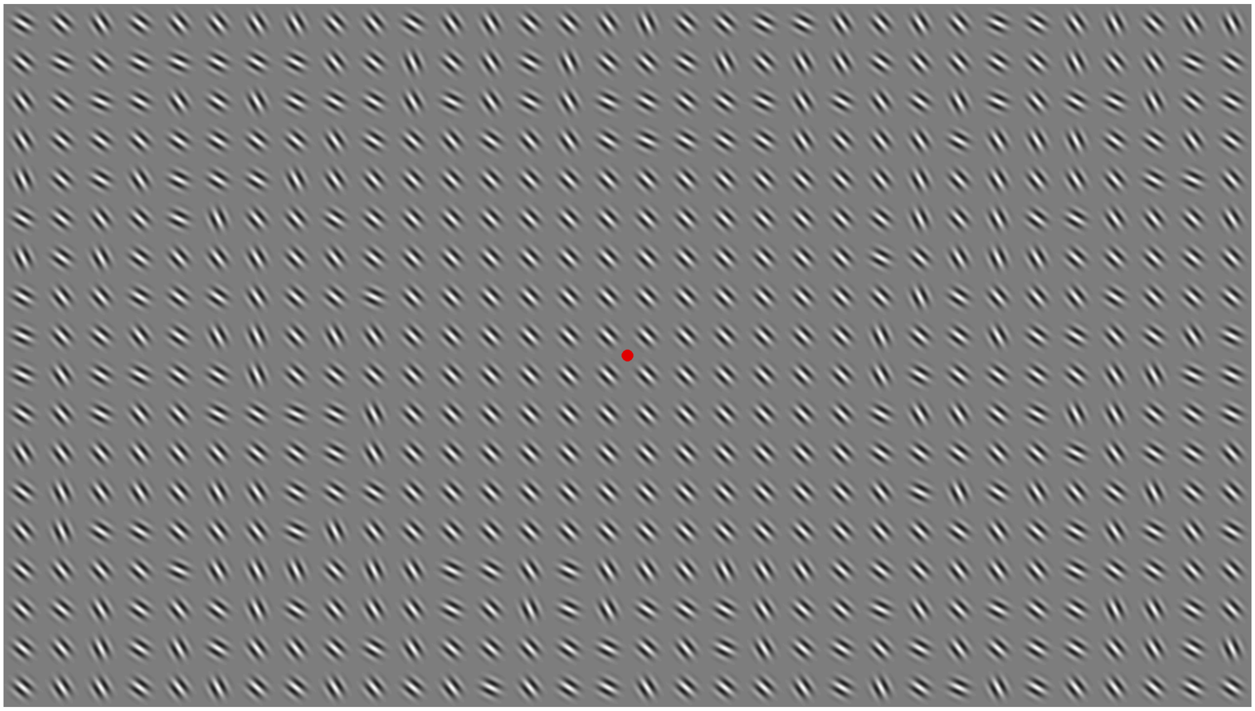

***Supplementary Figure 1.*** *The realistically scaled actual stimuli (with a matrix of 32 × 18 Gabors) used in the experiment. The central area (12 × 10 Gabors) has a coherent orientation (45 degrees clockwise or counterclockwise), while the peripheral area has randomly titled orientations (25 to 65 degrees). A central fixation dot maintains participants' gaze.*

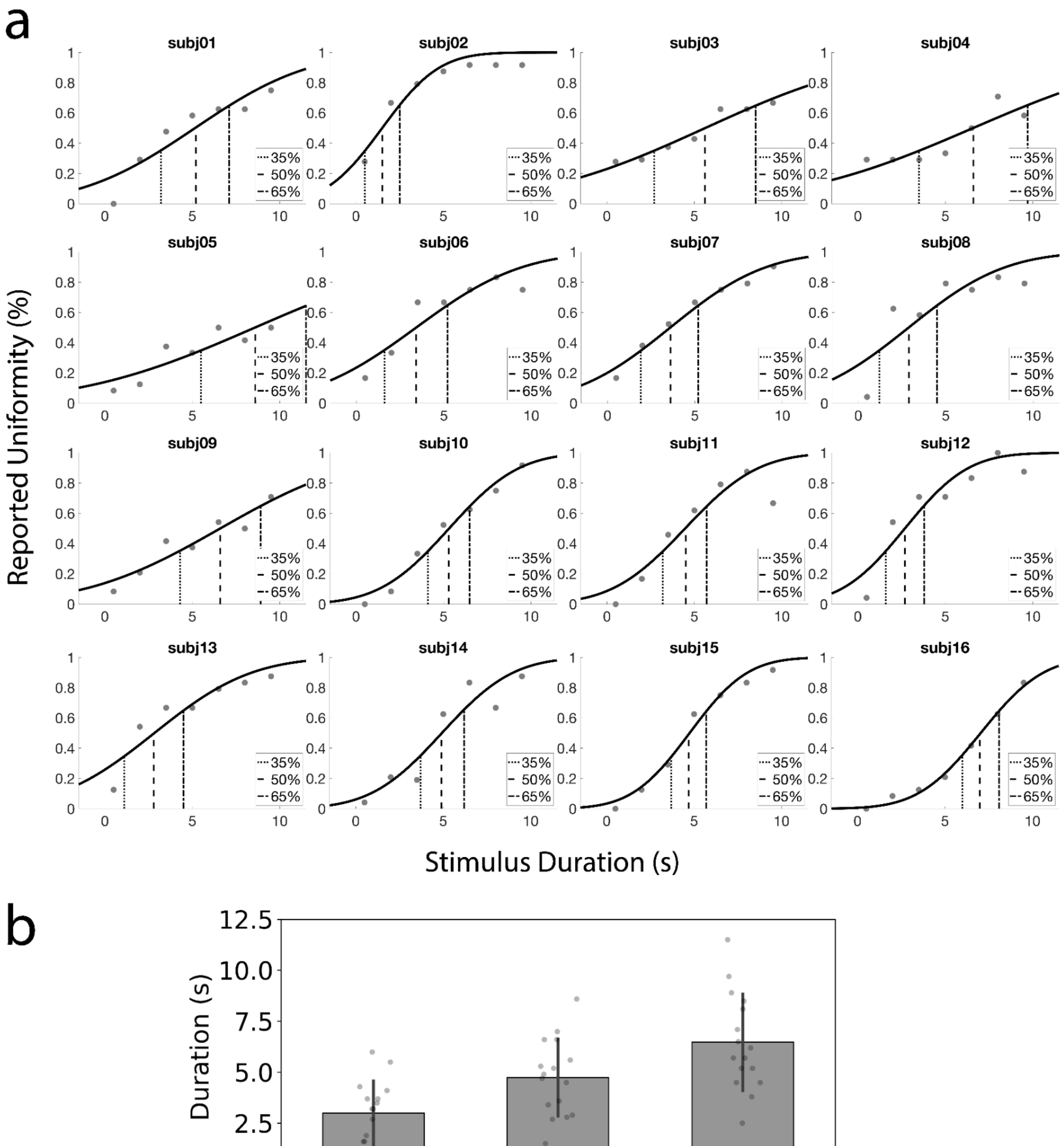


***Supplementary Figure 2.*** *Threshold duration measurements for each participant and group-level results.* ***a.*** *the fitted psychometric functions for all the participants (N=16), showing their 35%, 50%, and 65% threshold duration for the uniformity illusion.* ***b.*** *the group mean of durations for the 35%, 50%, and 65% conditions, with individual participant data represented as scatter plots. Error bars represent standard deviation.*

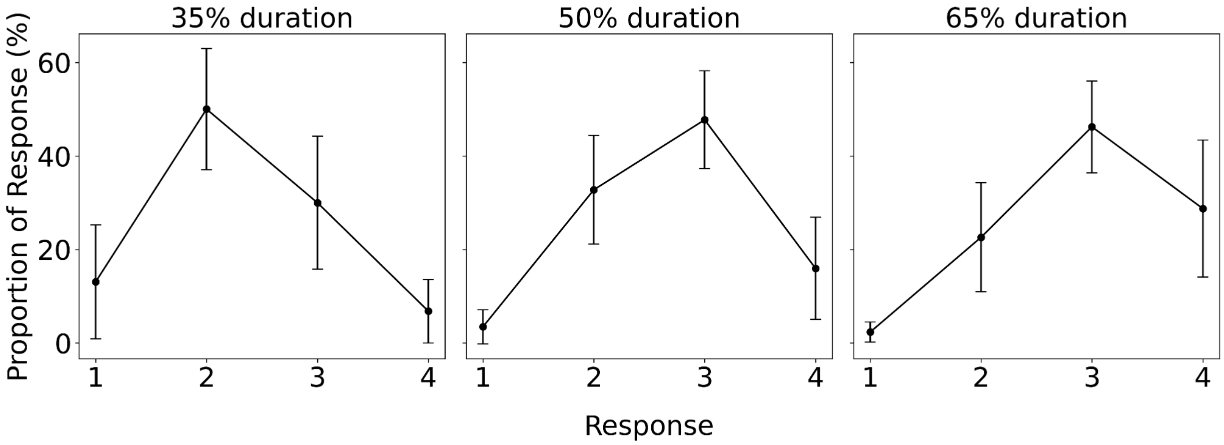


***Supplementary Figure 3.*** *Response distribution across threshold durations in the uniformity illusion task. This figure illustrates the normalized proportion of each of the four response categories across three different stimulus threshold durations: 35%, 50% and 65%. Each panel corresponds to one threshold duration, depicting how responses are distributed within that condition.*

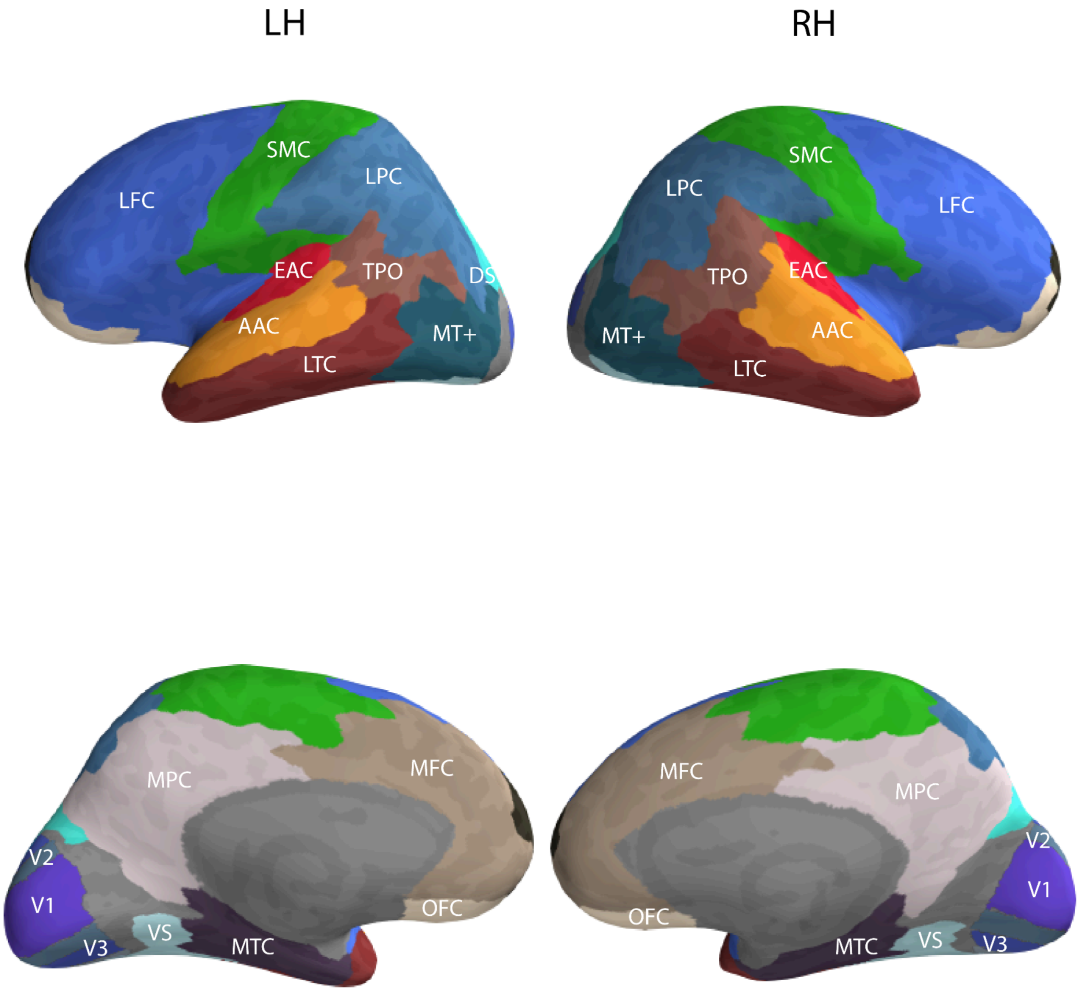


***Supplementary Figure 4.*** *Visualization of the Regions of Interest (ROIs) used in the study across the whole brain for one participant. The figure illustrates the ROIs employed in our study. The ROIs are defined based on the HCP-MMP atlas (Glasser et al., 2016) and are labeled as follows: LFC - lateral frontal cortex; MFC - medial frontal cortex; OFC - orbital frontal cortex; LPC - lateral parietal cortex; MPC - medial parietal cortex; SMC - sensori-motor cortex; TPO - temporal-parietal-occipital junction; LTC - lateral temporal cortex; MTC - medial temporal cortex; DS - dorsal stream; VS - ventral stream. For detailed mapping of our defined ROIs to the specific regions in the HCP-MMP atlas, please refer to Supplementary Table.1. V1 to V4 are delineated using a retinotopic mapping procedure.*

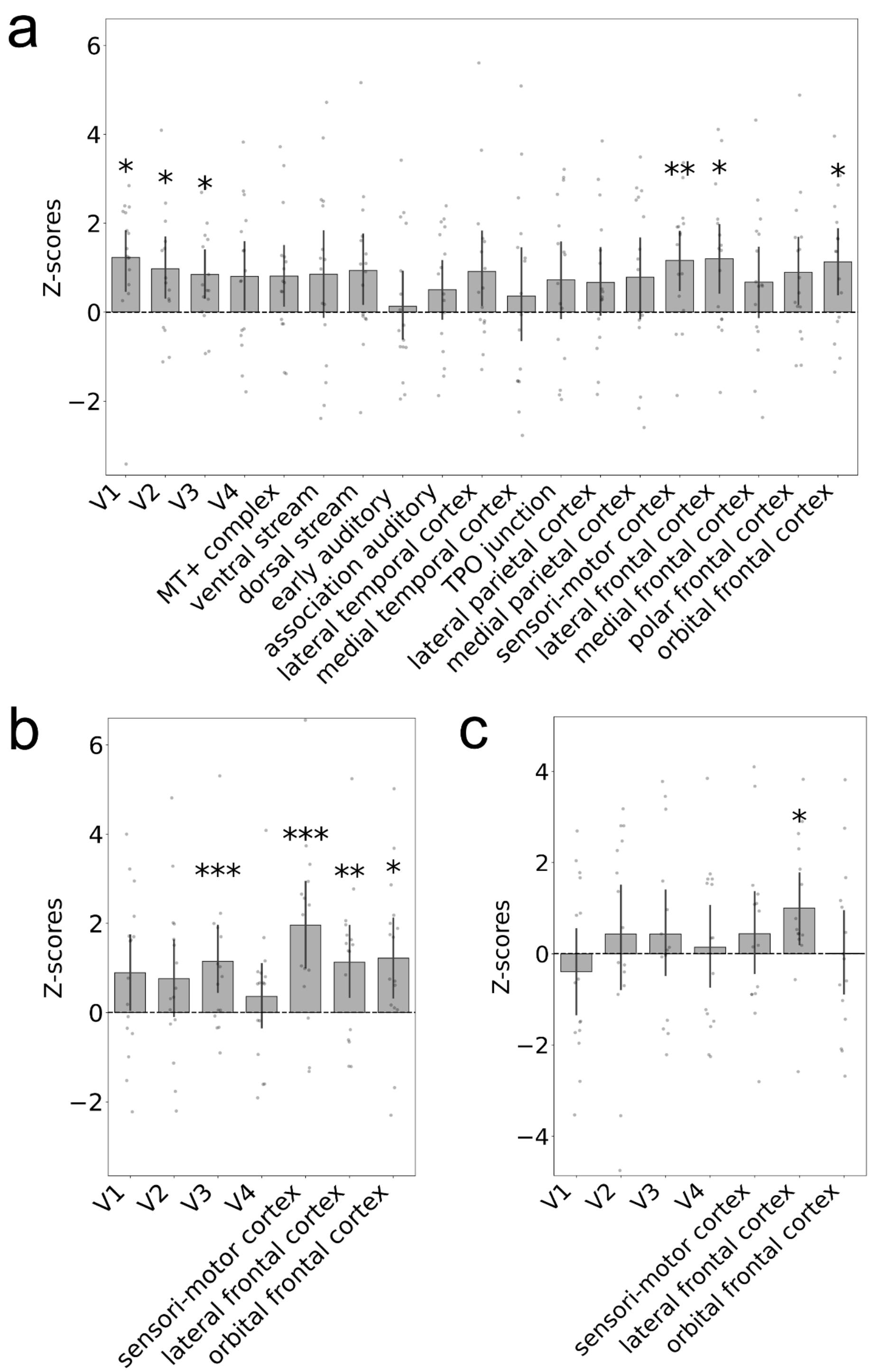


***Supplementary Figure 5.*** *Decoding and generalization results with permutation tests shuffling labels in both training and test sets, yielding similar outcomes.* ***a.*** *Decoding performance trained and tested on the 50% duration trials;* ***b.*** *generalization test performance on the 35% and 65% duration trials;* ***c.*** *generalization test performance on the movie frames. Error bars indicate 95% confidence interval obtained by bootstrap. * P<0.05, ** P<0.01, *** P<0.005 after correction.*

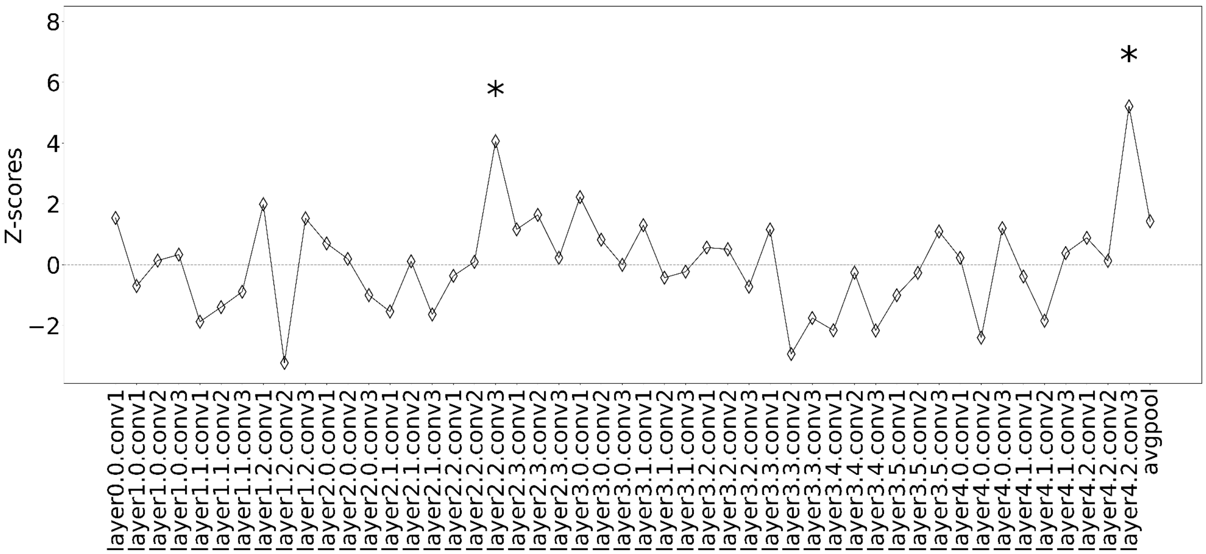


***Supplementary Figure 6.*** *decoding results on the features extracted from ResNet50 layers with permutation tests shuffling labels in both training and test set, which yielded similar outcomes. * P<0.05 after correction.*

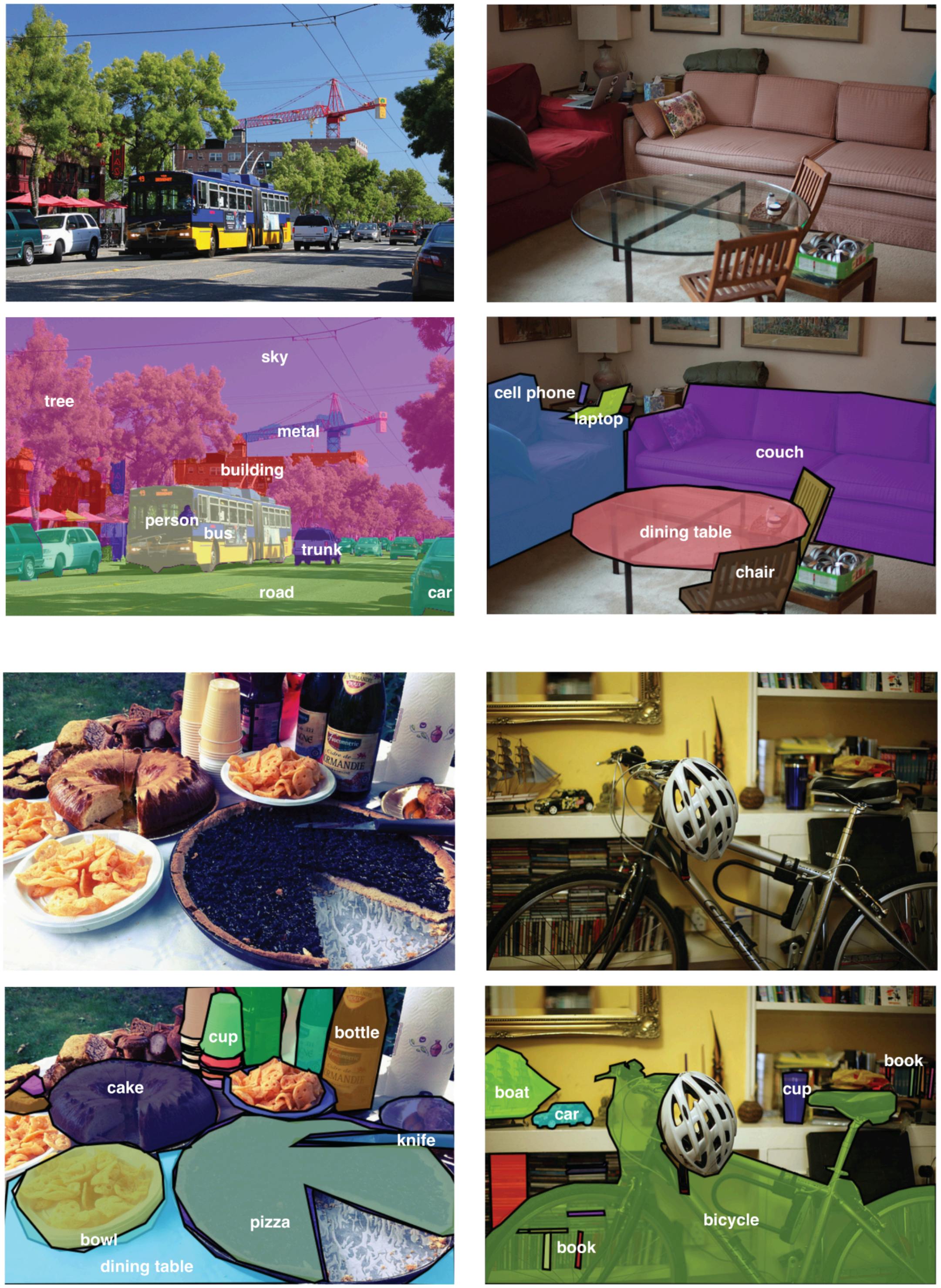


***Supplementary Figure 7.*** *Example figures with the semantic annotations in the COCO datasets. The different color patches in the lower panel represent different semantic categories.*

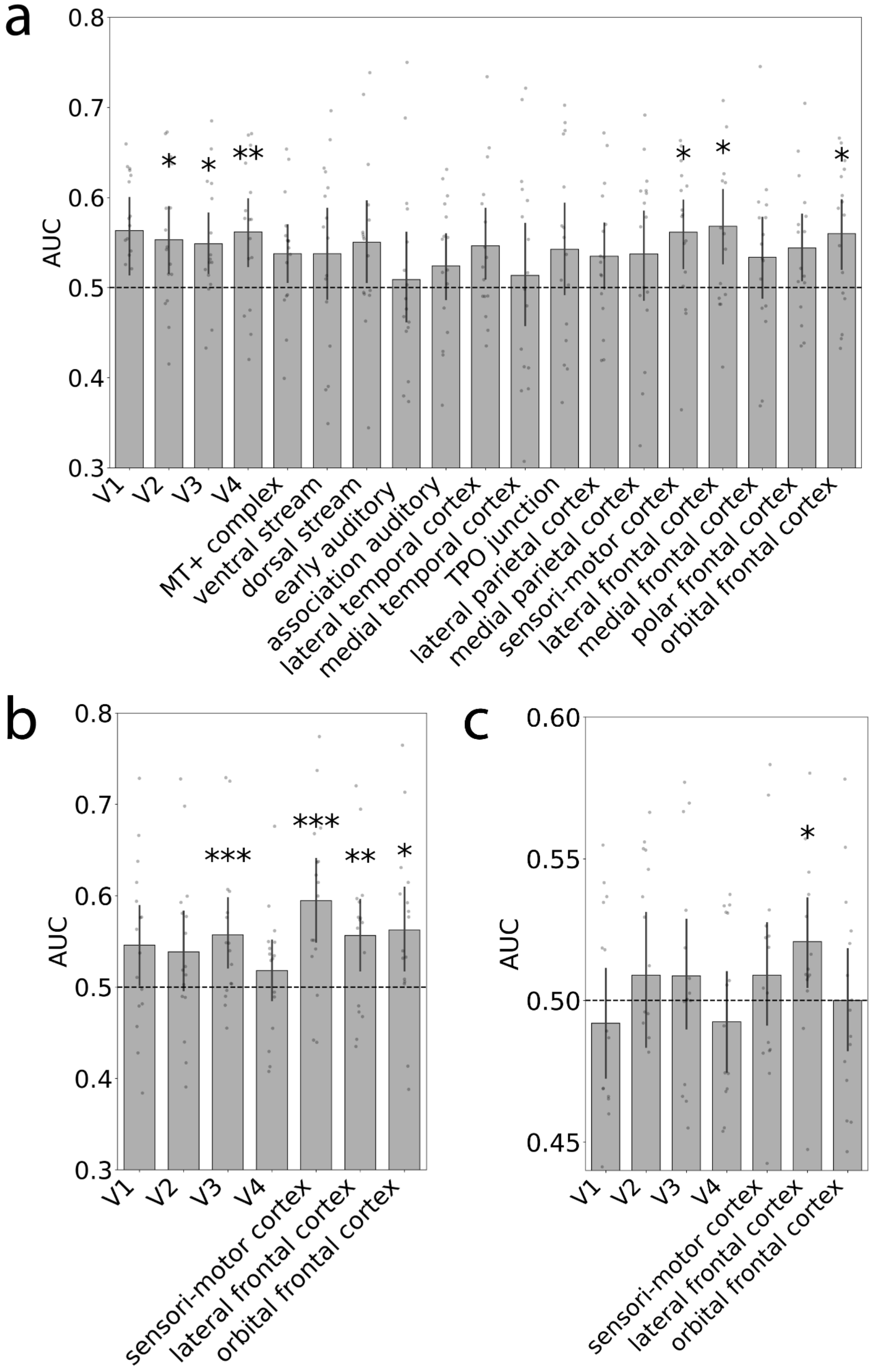


***Supplementary Figure 8.*** *Decoding and generalization results with raw AUC scores across participants, yielding similar results.* ***a.*** *Decoding performance trained and tested on the 50% duration trials;* ***b.*** *generalization test performance on the 35% and 65% duration trials;* ***c.*** *generalization test performance on the movie frames. Error bars indicate 95% confidence interval obtained by bootstrap. * P<0.05, ** P<0.01, *** P<0.005 after correction.*

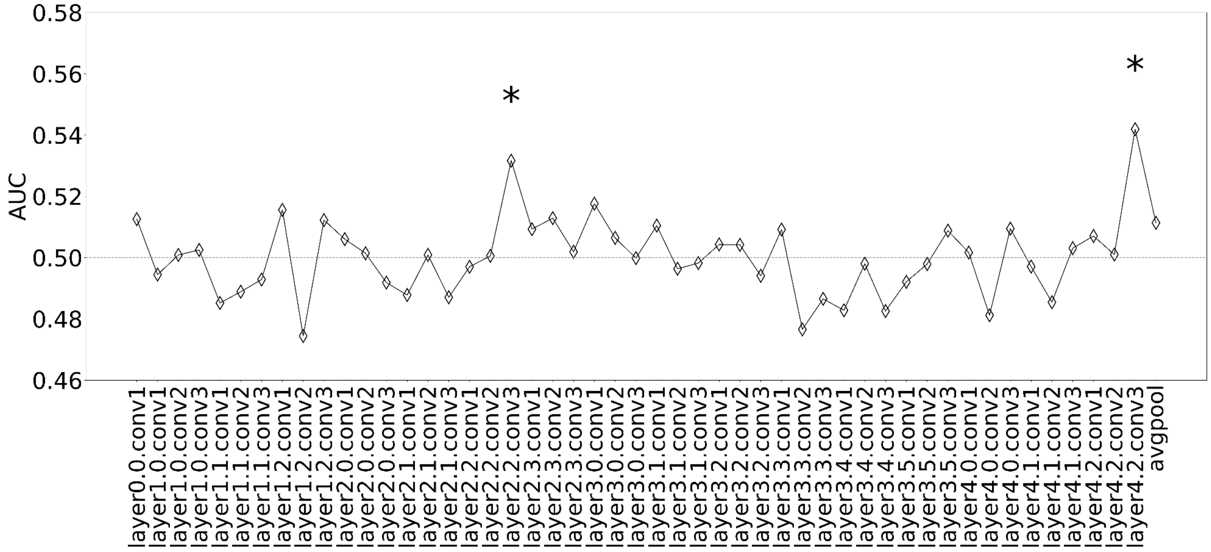


***Supplementary Figure 9.*** *Decoding results on the features extracted from ResNet50 layers with raw AUC scores. * P<0.05 after correction.*

| Defined ROIs | Corresponding HCP-MMP Atlas Regions |
|---|---|
| MT+ complex | MST, LO1, LO2, MT, PH, V4t, FST, V3CD, LO3 |
| Ventral stream | V8, FFC, PIT, VMV1, VMV3, VMV2, VVC |
| Dorsal stream | V6, V3A, V7, IPS1, V3B, V6A |
| Early auditory | A1, RI, PBelt, MBelt, LBelt |
| Association auditory | TA2, STGa, A5, STSda, STSdp, STSvp, A4, STSva |
| Lateral temporal cortex | TF, TGd, TE1a, TE1p, TE2a, TE2p, PHT, TGv, TE1m |
| Medial temporal cortex | EC, PreS, H, PeEc, PHA1, PHA3, PHA2 |
| TPO junction | PSL, STV, TPOJ1, TPOJ2, TPOJ3 |
| Lateral parietal cortex | 7Pm, 7AL, 7Am, 7PL, 7PC, LIPv, VIP, MIP, LIPd, AIP, PFt, PGp, IP2, IP1, IP0, PFop, PF, PFm, PGi, PGs |
| Medial parietal cortex | 23c, ProS, RSC, POS2, PCV, 7m, POS1, 23d, v23ab, d23ab, 31pv, DVT, 31pd, 31a |
| Sensori-motor cortex | 4, 3b, 1, 2, 3a, 5m, 5mv, 5L, 24dd, 24dv, SCEF, 6ma, 6mp, 43, OP4, OP1, OP2-3, FOP1, PFcm |
| Lateral frontal cortex | FEF, EF, 55b, 6d, 6a, 6v, 6r, 52, PoI2, FOP4, MI, Pir, AVI, AAIC, FOP3, FOP2, PoI1, Ig, FOP5, PI, 44, 45, 47l, IFJa, IFJp, IFSp, IFSa, p47r, SFL, 8Av, 8Ad, 8BL, 9p, 8C, p9-46v, 46, a9-46v, 9-46d, 9a, i6-8, s6-8 |
| Medial frontal cortex | p24pr, 33pr, a24pr, p32pr, a24, d32, 8BM, p32, 10r, 9m, 10v, 25, s32, a32pr, p24 |
| Polar frontal cortex | 10d, a10p, 10pp, p10p |
| Orbital frontal cortex | pOFC, 47m, 11l, 13l, OFC, 47s, a47r |

***Supplementary Table 1.*** *Mapping of defined ROIs to HCP-MMP Atlas Regions. This table lists the ROIs defined in our study and their corresponding labels from the HCP-MMP atlas (Glasser et al. 2016). The left column shows the ROIs used in our analyses, while the right column details the specific HCP-MMP atlas regions encompassed within each larger ROI.*

| ROI Name | Left Hemi (Mean %) | Left Hemi (SD %) | Right Hemi (Mean %) | Right Hemi (SD %) |
|---|---|---|---|---|
| 9-46d | 2.50 | 1.38 | 2.81 | 2.04 |
| 8Ad | 2.40 | 1.93 | 1.85 | 0.87 |
| 6a | 2.30 | 1.92 | 2.82 | 1.89 |
| 6r | 2.26 | 1.66 | 2.68 | 2.06 |
| 8C | 2.20 | 1.61 | 1.75 | 1.02 |
| IFSa | 2.19 | 1.35 | 1.47 | 0.62 |
| p47r | 2.09 | 1.39 | 1.15 | 1.35 |
| p9-46v | 1.91 | 1.78 | 1.81 | 1.47 |
| 8Av | 1.80 | 1.41 | 1.23 | 0.84 |
| a9-46v | 1.67 | 1.28 | 0.90 | 0.89 |
| 9a | 1.59 | 1.50 | 1.73 | 1.91 |
| 45 | 1.58 | 1.16 | 1.09 | 0.81 |
| SFL | 1.57 | 0.91 | 0.57 | 0.64 |
| 46 | 1.46 | 0.69 | 2.31 | 1.12 |
| 6d | 1.35 | 1.50 | 1.08 | 0.78 |
| 8BL | 1.23 | 0.99 | 1.52 | 1.81 |
| 47l | 1.23 | 1.39 | 1.24 | 0.88 |
| AVI | 1.21 | 1.07 | 1.64 | 1.42 |
| FEF | 1.11 | 1.27 | 1.90 | 1.42 |
| IFJa | 1.05 | 1.17 | 0.66 | 0.78 |
| PoI1 | 1.03 | 0.84 | 0.89 | 0.84 |
| FOP4 | 0.97 | 0.78 | 0.84 | 0.91 |
| 9p | 0.95 | 0.86 | 1.34 | 1.62 |
| 44 | 0.95 | 0.78 | 1.29 | 0.99 |
| IFSp | 0.92 | 0.66 | 1.39 | 1.04 |
| s6-8 | 0.92 | 0.95 | 0.92 | 0.82 |
| PoI2 | 0.84 | 0.91 | 1.19 | 1.36 |
| FOP5 | 0.82 | 0.63 | 1.37 | 1.76 |
| MI | 0.82 | 0.70 | 0.75 | 0.49 |
| 52 | 0.78 | 0.96 | 0.35 | 0.42 |

| | | | | |
|---|---|---|---|---|
| FOP3 | 0.65 | 0.73 | 0.27 | 0.47 |
| AAIC | 0.61 | 0.61 | 0.65 | 0.83 |
| Ig | 0.58 | 0.56 | 0.83 | 0.92 |
| 55b | 0.57 | 0.53 | 1.11 | 1.25 |
| Pir | 0.55 | 0.44 | 0.92 | 0.66 |
| 6v | 0.53 | 0.41 | 1.27 | 0.99 |
| i6-8 | 0.53 | 0.35 | 0.78 | 0.68 |
| PI | 0.50 | 0.69 | 0.70 | 0.86 |
| IFJp | 0.50 | 0.50 | 0.67 | 0.70 |
| FOP2 | 0.45 | 0.53 | 0.23 | 0.35 |
| EF | 0.26 | 0.29 | 0.57 | 0.67 |

***Supplementary Table 2.*** *Statistical summaries for the decoding weights in the lateral frontal cortex across participants for the uniformity illusion data. The anatomical ROIs are taken from HCP-MMP atlas (Glasser et al. 2016). There are 41 subregions in the lateral frontal cortex. The table shows the group mean and standard deviation of the percentage of the selected 1200 vertices within each subregion in the left and right hemispheres across participants.*
